# *Drosophila* menthol sensitivity and the Precambrian origins of TRP-dependent chemosensation

**DOI:** 10.1101/690933

**Authors:** Nathaniel J. Himmel, Jamin M. Letcher, Akira Sakurai, Thomas R. Gray, Maggie N. Benson, Daniel N. Cox

## Abstract

Transient receptor potential (TRP) cation channels are highly conserved, polymodal sensors which respond to a wide variety of stimuli. Perhaps most notably, TRP channels serve critical functions in nociception and pain. A growing body of evidence suggests that TRPM (<u>M</u>elastatin) and TRPA (<u>A</u>nkyrin) thermal and electrophile sensitivities predate the protostome-deuterostome split (>550 million years ago). However, TRPM and TRPA channels are also thought to detect modified terpenes (*e.g*., menthol). Although terpenoids like menthol are thought to be aversive and/or harmful to insects, mechanistic sensitivity studies have been largely restricted to chordates. Furthermore, it is unknown if TRP-menthol sensing is as ancient as thermal and/or electrophile sensitivity. Combining genetic, optical, electrophysiological, behavioural, and phylogenetic approaches, we tested the hypothesis that insect TRP channels play a conserved role in menthol sensing. We found that topical application of menthol to *Drosophila melanogaster* larvae elicits a *Trpm*- and *TrpA1*-dependent nocifensive rolling behaviour, which requires activation of Class IV nociceptor neurons. Further, in characterizing the evolution of TRP channels, we put forth the hypotheses that 3 previously undescribed TRPM channel clades (basal, αTRPM, and βTRPM), as well as TRPs with residues critical for menthol sensing, were present in ancestral bilaterians.

## Introduction

Menthol and icilin are often referred to as “cooling agents” – while these chemicals do not physically chill, topical application typically elicits a pleasant cooling sensation in humans [1]. Perhaps owing to its perceived cooling properties, menthol has been used as a topical analgesic, often to reduce the severity of itching and/or burning sensations [2–4]. That said, what is pleasant, harmful, or potentially painful will often be species-dependent. It has been previously reported that menthol affects the behaviour of insects; for example, menthol-infused foods are aversive to *Drosophila melanogaster*, and there is some evidence that menthol functions as an insecticide [5–11]. However, relatively little is known concerning the mechanisms by which insects sense and respond to the cooling agents, as previous studies have focused largely on deuterostomes, and, with respect to molecular determinants among those species, chiefly on terrestrial chordates [12–20]. By extension, it is unknown if possible shared mechanisms have their origins in a common ancestor.

Like many other compounds, menthol and icilin are thought to be detected by TRP channels – primarily TRPM8 and TRPA1 [17, 21, 22]. TRP channels are variably selective cation channels which are differentially gated by a wide variety of thermal, chemical, and mechanical stimuli. It is the multimodal nature of TRPM8 and TRPA1 – in humans constituted by at least menthol, icilin, and cold sensing – which partly underlies the similar phenomenological character (*i.e*., cooling) associated with these very different stimuli [22–24].

The chordate TRPM (<u>TRP</u> <u>M</u>elastatin) family is typically divided into 8 distinct paralogues (TRPM1-TRPM8) thought to have emerged sometime prior to the divergence of tetrapods and fish (although fish are thought to have lost their TRPM8 orthologue) [25]. TRPM8 is multimodal, responding to cold, menthol, and in mammals, icilin [22, 26]. It has been suggested that menthol directly binds with the TRPM8 Voltage Sensor-Like Domain (VSLD), and gating requires interactions between this domain, bound menthol molecules, and the highly conserved C-terminal TRP domain [13, 20]. TRPM8-menthol gating has been well characterized in mammalian channels, and a number of critical amino acid residues have been identified in both the VSLD and the TRP domain [12, 14, 16, 19, 20].

In contrast to the TRPM family, the chordate TRPA (<u>TRP</u> <u>A</u>nkyrin) family is very small, typically containing only a single member, TRPA1 [27, 28]. TRPA1 is a polymodal nociceptor involved in the detection of noxious cold, noxious heat, menthol, icilin, and electrophilic chemicals such as allyl isothiocyanate (AITC, found in mustard oil and wasabi) [21, 24, 29, 30]. It has been suggested that TRPA1 menthol sensitivity is linked to specific serine and threonine residues found in transmembrane segment 5, and that several nearby residues are responsible for species-specific TRPA1-menthol interactions [31].

Insects lack a true TRPM8 orthologue. In fact, the *Drosophila* genomes contains only a single TRPM family gene, *Trpm* (**Figure 1, top**) [32]. While separated by several gene duplication events and more than 550 million years of evolution, *Drosophila* Trpm and its chordate counterpart, TRPM8, are both involved in cold sensing [33]. With respect to TRPAs, the *Drosophila* genome encodes 4 TRPA family genes (**Figure 1, bottom**): *TrpA1* (the homologue to chordate *TRPA1*), *painless* (*pain*), *water witch* (*wtrw*), and *pyrexia* (*pyx*). Like vertebrate TRPA1, *Drosophila* TrpA1 has been implicated in high-temperature and chemical nociception [27, 33–37].

**Figure 1.**
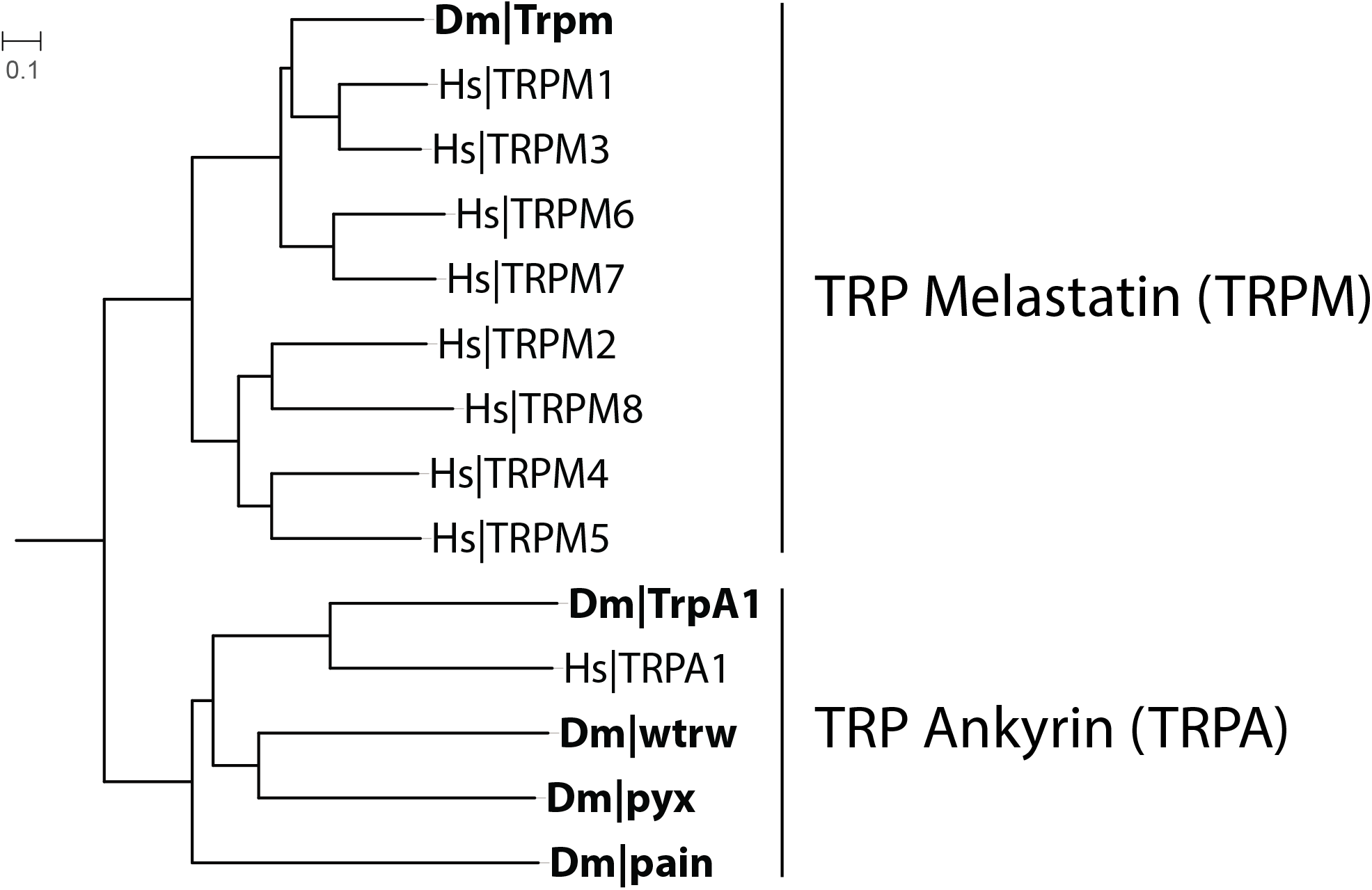
The *Drosophila melanogaster* (Dm) genome encodes 1 TRPM (Trpm) and 4 TRPA (TrpA1, painless, pyrexia, and water witch) channels. Mid-point rooted tree of *Drosophila* (Dm) and human (Hs) TRPMs and TRPAs.

TRPA1 high-temperature and electrophile sensitivities are present in both protostomes and deuterostomes [27, 36–45]. While current evidence suggests – despite some controversy – that TRPA1 cold sensitivity evolved relatively recently, perhaps among early synapsids (<300 mya) [46–48], TRPM cold sensitivity is conserved in insects [33]. As such, a growing body of evidence suggests that TRP channel thermal and electrophile sensitivity may have a common origin predating the protostome-deuterostome split (>550 mya). However, it remains unknown if TRPA- and TRPM-dependent menthol sensitivity is equally ancient.

In order to further elucidate the evolutionary history of TRPM and TRPA channels, we explored the mechanisms by which *Drosophila* senses and responds to cooling agents. Herein, we report that topical menthol application elicits a dose-dependent nocifensive rolling behaviour in *Drosophila*. Further, we use a suite of genetic, optical, electrophysiological, and behavioural approaches to demonstrate that the menthol-evoked response is *Trpm*- and *TrpA1*-dependent, and acts via Class IV (CIV) nociceptors. Finally, in light of these findings, we assess the evolutionary histories and sequence homologies of TRPM and TRPA channels. In doing so, we characterize several putative TRPM and TRPA channels in *Acropora digitifera* (coral), *Strongylocentrotus purpuratus* (purple sea urchin), *Octopus bimaculoides* (California two-spot octopus), *Priapulus caudatus* (penis worm), and *Aplysia californica* (California sea hare), as well as outline the discovery of three previously undescribed TRPM channel clades (basal TRPMs, αTRPMs, and βTRPMs), which we hypothesize were present in the last common bilaterian ancestor, Urbilateria. The results of these studies add to a growing body of evidence suggesting that chemosensation and nociception have functional origins in the Precambrian world [49].

## Materials & Methods

### Animals

All *Drosophila melanogaster* stocks were maintained at 24°C under a 12:12 light:dark cycle. In order to accelerate development, genetic crosses (and relevant controls) were raised for 5 days at 29°C under a 12:12 light:dark cycle. Wandering 3rd instar larvae were used for all experiments. The OregonR (ORR) strain was used to assess wild-type behaviour, and, where appropriate, the *w^1118^* strain and outcrossed parental strains were used as genetic background controls. Transgenic and mutant strains included: *Trpm^2^* and *wtrw^1^* (gifts of K. Venkatachalam); *TrpA1^W903*^* and *TrpA1^1^* (gifts of W. D. Tracey); *pyx^3^, pain^70^*, and *Trpm* deficiency *Df(2R)XTE-11* (gifts of M. J. Galko); *GAL4^GMR57C10^* (pan-neuronal driver, BDSC #39171); *GAL4^ppk^* (Class IV driver, BDSC #32079); *UAS-CaMPARI* (BDSC #58763); *UAS-TeTxLC* (active tetanus toxin, BDSC #28837); *UAS-IMP-TNT^VI-A^* (inactive tetanus toxin, BDSC #28840); *UAS-TrpA1-RNAi* 1 (BDSC #31384); *UAS-TrpA1-RNAi* 2 (BDSC #66905); *UAS-Trpm-RNAi* 1 (BDSC #31672); and *UAS-Trpm-RNAi* 2 (BDSC #31291).

### Behaviour

L(-)-menthol (Acros Organics) and icilin (Tocris Bioscience) were suspended in liquid DMSO at various concentrations. Solutions were discarded after 24 hours. Individual larval subjects were placed in a well of a glass 9-well plate, and 10μL of solution was delivered to the well via micropipette. Subjects were observed for 60 seconds and their behaviours recorded. In order to increase subject-background contrast in the representative images/videos, animals were left to freely locomote on a moist black arena, and menthol was applied as above.

### CaMPARI-based Ca^2+^ analyses

*UAS-CaMPARI* expression was driven via *GAL4^GMR57C10^*. Live, freely behaving larvae were exposed to 8% menthol or vehicle as described above. Photoconverting light was delivered under a Zeis AxioZoom V16 microscope as previously described [50, 51]. Larvae were subsequently placed in 1 drop of 1:5 diethyl ether:halocarbon oil and secured between a slide and slide cover. Class IV neurons were imaged via a Zeiss LSM780 confocal microscope, and the resulting z-stacks were volume rendered as two-dimensional maximum intensity projections. Red and green fluorescence intensity was assessed using the FIJI distribution of ImageJ software.

### Electrophysiology

To record single-unit spiking activity of Class IV neurons, we first prepared fillet preparations from *GAL4^ppk^>UAS-mCD8::GFP* larvae. Larvae were placed in a petri dish lined with Sylgard^®^ 184 (Dow Corning, Midland, MI) filled with HL-3 saline. The ventral body wall was cut open with fine scissors, and all muscles were carefully removed with a polished tungsten needle and scissors. The preparation was then constantly superfused with HL-3 saline at a rate of 1 mL/min at room temperature, and allowed to rest for more than 1 hour before recording. Menthol was first dissolved in DMSO at a concentration of 500 mM and then diluted in HL-3 saline to a final concentration 0.5 mM. Menthol was bath-applied to the specimen through superfusion. Extracellular recordings were made with a macropatch pipette (tip diameter = 5-10 μm) connected to the headstage of a patch-clamp amplifier (AxoPatch200B, Molecular Devices, Sunnyvale, CA). Gentle suction was applied to draw the soma and a small portion of neurite into the pipette. The amplifier was set to the track mode to record neuronal spikes. The output signals from the amplifier were digitized at a sampling frequency of 10 kHz using a Micro1401 A/D converter (Cambridge Electronic Design, Cambridge, UK) and acquired into a laptop computer running Windows 10 with Spike2 software ver. 8 (Cambridge Electric Design, Cambridge, UK). Average spike frequency was measured in a 30 sec time window during baseline control conditions, superfusion of vehicle, and superfusion of menthol.

### Phylogenetics

Amino acid sequences for previously characterized TRP channels were collected from the following databases: JGI Genome (*Monosiga brevicollis, Nematostella vectensis*, and *Daphnia pulex*), Ensembl (*Danio rerio, Gallus gallus*, and *Strigamia maritima*), NCBI (*Hydra vulgaris, Homo sapiens, Mus musculus, Apis mellifera*, and *Galendromus occidentalis*), WormBase (*Caenorhabditis elegans*), or FlyBase (*Drosophila melanogaster*). *Panulirus argus* sequences were collected from [52]. In order to identify novel TRPM and TRPA channels, publically available protein models based on genomic and/or transcriptomic sequences for *Acropora digitifera* [53], *Strongylocentrotus purpuratus* [54, 55], *Octopus bimaculoides* [56], *Priapulus caudatus* (BioProject PRJNA20497, GenBank AXZU00000000.2), and *Aplysia californica* (BioProject, PRJNA209509, GenBank AASC00000000.3) were pBLASTed [57] against *D. melanogaster* TRP sequences. Sequences >200aa in length and with an E-value <1E-30 were retained and subsequently analyzed via InterProScan [58]. Sequences which had predicted transmembrane segments and characteristic Ankyrin repeats (for TRPAs) were retained. Accession numbers are available in the supplementary material (**Table S1**).

Amino acid sequences were MUSCLE [59] aligned in MEGA7 [60]. Poorly aligned regions and spurious sequences were identified and trimmed using automated methods packaged with TrimAl [61]. As TRPA Ankyrin repeats were generally poorly aligned, they were manually removed prior to automated trimming by excluding everything N-terminal of 20aa before the start of predicted transmembrane segment 1. IQ-Tree was then used to perform a composition chi^2^ test in order to assess sequence homogeneity within TRP subfamilies [62]. Due to extreme divergence, and in order to minimize possible topology disruptions due to long branch attraction [63–65], *P. argus* TRPMm, TRPA-like1, and TRPA5-like1, and *C. elegans* TRPA-2 were excluded from final analyses; each failed the composition chi^2^ test and introduced extremely long branches with weak support in a first-pass analysis. Final, trimmed alignments were used to generate phylogenetic trees. Bayesian trees were constructed in MrBayes (version 3.2.6) using a mixed amino acid substitution model and a gamma distributed rate of variation [66, 67]. Two independent MCMC analyses (initial settings: 1,000,000 chains with sampling every 10) were run until convergence (<0.05). 25% of the chain was considered burn-in and discarded. Maximum likelihood trees were constructed in IQ-Tree using an amino acid substitution model automatically selected by ModelFinder [68]. Ultrafast bootstrapping (2000 bootstraps) was also performed in IQ-Tree [69]. Trees were visualized and edited in iTOL [70] and Adobe Illustrator CS6. Branches with low support (posterior probability <0.7 or bootstrap <70) were considered unresolved, and were collapsed to polytomies in the final trees. Ancestral sequence predictions were made in MEGA7 using the maximum likelihood approach against previously generated alignments and Bayesian trees [71].

### Statistical analyses

Population proportions are presented as % ± standard error of the proportion (SEP); differences in proportion were assessed using a generalized linear model with a logit link and a binomial error distribution, and pairwise comparisons were made using the lsmeans R package [72] and the Tukey method for multiple comparisons. CaMPARI photoconversion, spike frequency, and latency measures are presented as mean ± standard error of the mean (SEM); differences were assessed in GraphPad PRISM (GraphPad Software, La Jolla, California, USA) by unpaired t-test or ANOVA with Dunnett’s *post-hoc* test. Asterisk (*) indicates statistical significance p<0.05, with significant p-values (as compared to wild-type or appropriate control) listed in the associated figure legend.

## Results

### Menthol elicits *Trpm*- and *TrpA1*-dependent nocifensive rolling in *Drosophila* larvae

Vehicle (DMSO), menthol, or icilin was topically applied to freely behaving *Drosophila melanogaster* larvae, and their behaviour recorded (10μL, delivered via micropipette). Mentholated solutions elicited a stereotyped rolling behavior, identical to a previously described [73, 74] nocifensive response (**Figure 2A; Movie S1**). The proportion of animals which responded to menthol increased with higher menthol concentrations (**Figure 2B**). In contrast, treatment with DMSO (**Figure 2B, vehicle; Movie S2**) or icilin (**Figure 2B; Movie S3**) did not elicit rolling.

**Figure 2.**
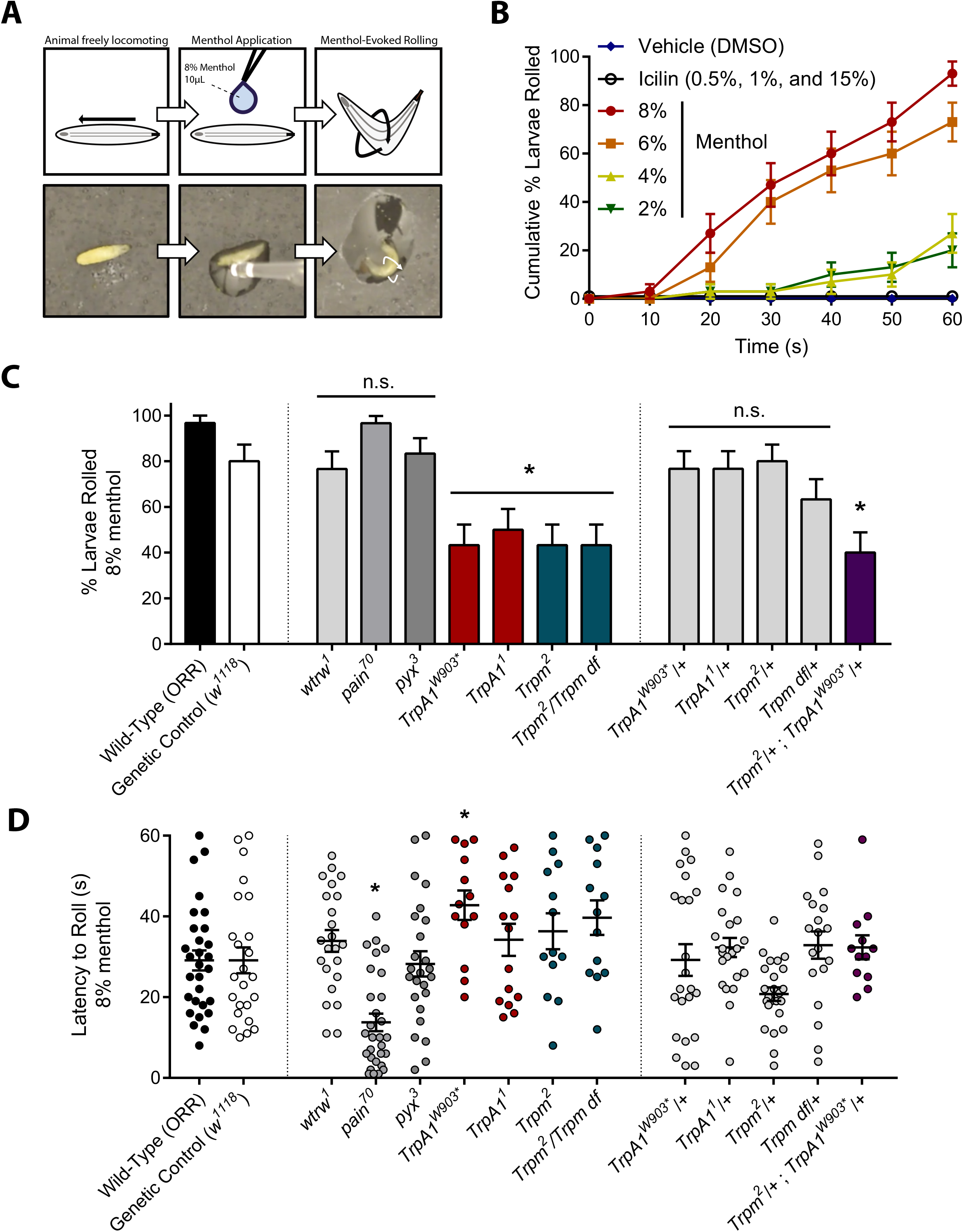
*Drosophila* larvae exhibit nocifensive rolling in response to menthol. **(A)** Menthol-evoked rolling. Top, cartoon. Bottom, video stills. **(B)** Cumulative proportion of wild-type rollers in response to menthol. The rolling response increased at higher menthol concentrations. Cumulative proportion ± SEP. n=30 for each condition. **(C)** Proportion of rollers in response to 8% menthol. Compared to wild-type, fewer *TrpA1* and *Trpm* mutants rolled in response to menthol (*TrpA1^W903*^*, p=0.0221; *TrpA1*^1^ p=0.0479; *Trpm*^2^, p=0.0221; *Trpm*^2^/Trpm df, p=0.0221). *Trpm* and *TrpA1* are haplosufficient for menthol sensitivity, but fewer transheterozygous mutants rolled in response to menthol (p=0.0090). Proportions represented as % rollers ± SEP. n=30 for each condition. **(D)** Latency to roll in response to 8% menthol. *painless* mutation sensitized rollers (p=0.0006), while *TrpA1^W903*^* mutation desensitized rollers (p=0.0388). Latency represented as time to roll in seconds ± SEM. n proportional to % rollers, where initial sample size was 30.

Given TRP-dependent mechanisms of menthol sensing in vertebrates, we hypothesized that menthol-evoked rolling requires TRPM and TRPA channels. To test this, we assessed the behaviour of whole-animal *Trpm, TrpA1, pyx, pain*, and *wtrw* mutants. Compared to control, a significantly smaller proportion of homozygous *TrpA1* and *Trpm* mutants rolled in response to menthol (**Figure 2C**). Additionally, we observed a significantly increased latency to roll for those *TrpA1^W903*^* mutants that did respond (**Figure 2D**). In contrast, *pain* mutation sensitized larvae to menthol – homozygous *pain* mutants rolled with severely decreased latency, and many rolled immediately following menthol application (**Figure 2D**). Given that *painless* mutants could be sensitized to the point of immediately rolling following menthol application, it appears likely that menthol was rapidly delivered across the cuticle.

As *TrpA1* and *Trpm* were both required for the menthol-evoked response, mutations were tested combinatorially. While heterozygous *TrpA1* (*TrpA1^W903*^*/+ and *TrpA1*^1^/+) and *Trpm* (*Trpm^2^*/+ and *Trpm* df/+) mutants had no significant behavioural defects, transheterozygous (*TrpA1*^W903*^/+;*Trpm*^2^/+) mutants exhibited significantly inhibited menthol-evoked rolling, indicating that these channels genetically interact in menthol sensing (**Figure 2C**).

### Class IV nociceptors mediate menthol-evoked rolling

*Drosophila* larvae have two known classes of peripheral nociceptors: multidendritic (md) Class III neurons are cold nociceptors and innocuous touch mechanosensors, whereas md Class IV (CIV) neurons are polymodal nociceptors which detect noxious heat, mechanical insults, and short-wavelength light [33, 34, 74–78]. As rolling behaviour has been previously associated with CIV nociceptor activation, we hypothesized that menthol-evoked rolling requires CIV activation. In order to assess activation patterns *in vivo*, the *GAL4-UAS* system was used to drive neuronal expression of CaMPARI [79], a genetically encoded Ca^2+^ indicator. Live, freely behaving transgenic animals were exposed to menthol or vehicle, as above. The level of CIV neural activation was assessed *post-hoc*, in live animals, as a green-to-red shift in CaMPARI fluorescence, which occurs in the presence of violet light as a function of increases in intracellular Ca^2+^ concentration. CaMPARI analyses revealed significant activation of CIV nociceptors in response to menthol, relative to vehicle control (**Figure 3A**). To directly assess menthol-evoked activation of CIV neurons, we performed single-unit electrophysiological recordings of CIV neurons. Electrophysiological recordings demonstrate that superfusion of menthol significantly increases spike frequency in CIV neurons relative to controls (**Figure 3B**).

**Figure 3.**
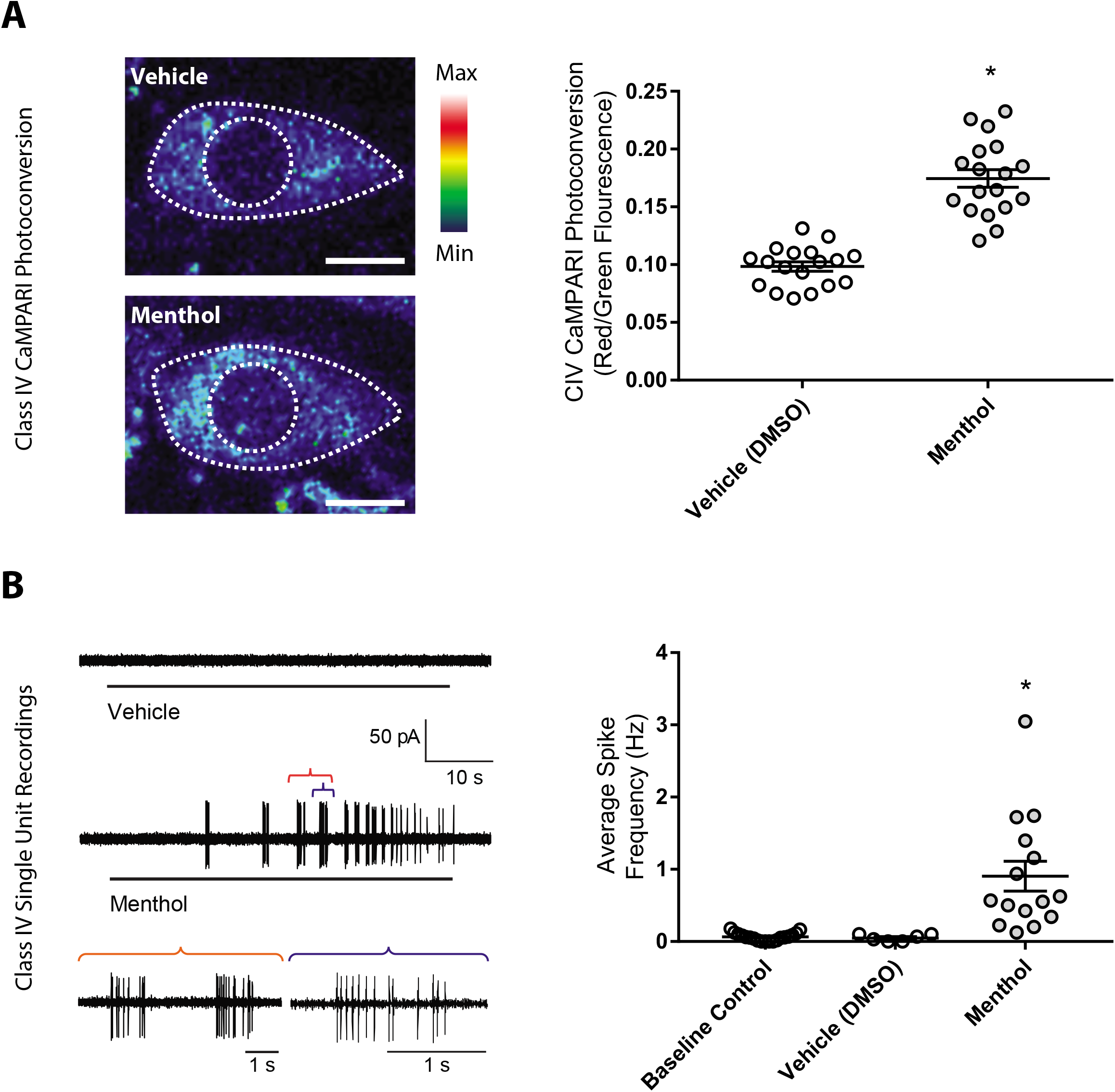
Menthol exposure activates Class IV (CIV) nociceptors. **(A)** (left) Representative CIV CaMPARI photoconversion visualized using a 16-color lookup table. (right) CIV neurons were significantly activated by menthol, as compared to control (p<0.0001). Quantification of CaMPARI photoconversion by F_red_/F_green_ CaMPARI fluorescence intensities. Photoconversion presented as mean ± SEM. n=18 for each condition. **(B)** (left) Representative CIV single-unit recordings, with insets showing menthol-evoked spiking activity. (right) Menthol superfusion elicited significantly increased average spike frequency in CIV neurons (p=0.0001). Average frequency in Hz ± SEM. Saline control, n=18; DMSO control, n=6; menthol, n=15.

In order to test the necessity of CIV neurons for menthol-evoked rolling, genetically encoded active tetanus toxin (TNT) was used to block synaptic transmission in CIV neurons. GAL4^ppk^-driven, CIV-specific expression of TNT completely ablated menthol-evoked rolling (**Figure 4**).

**Figure 4.**
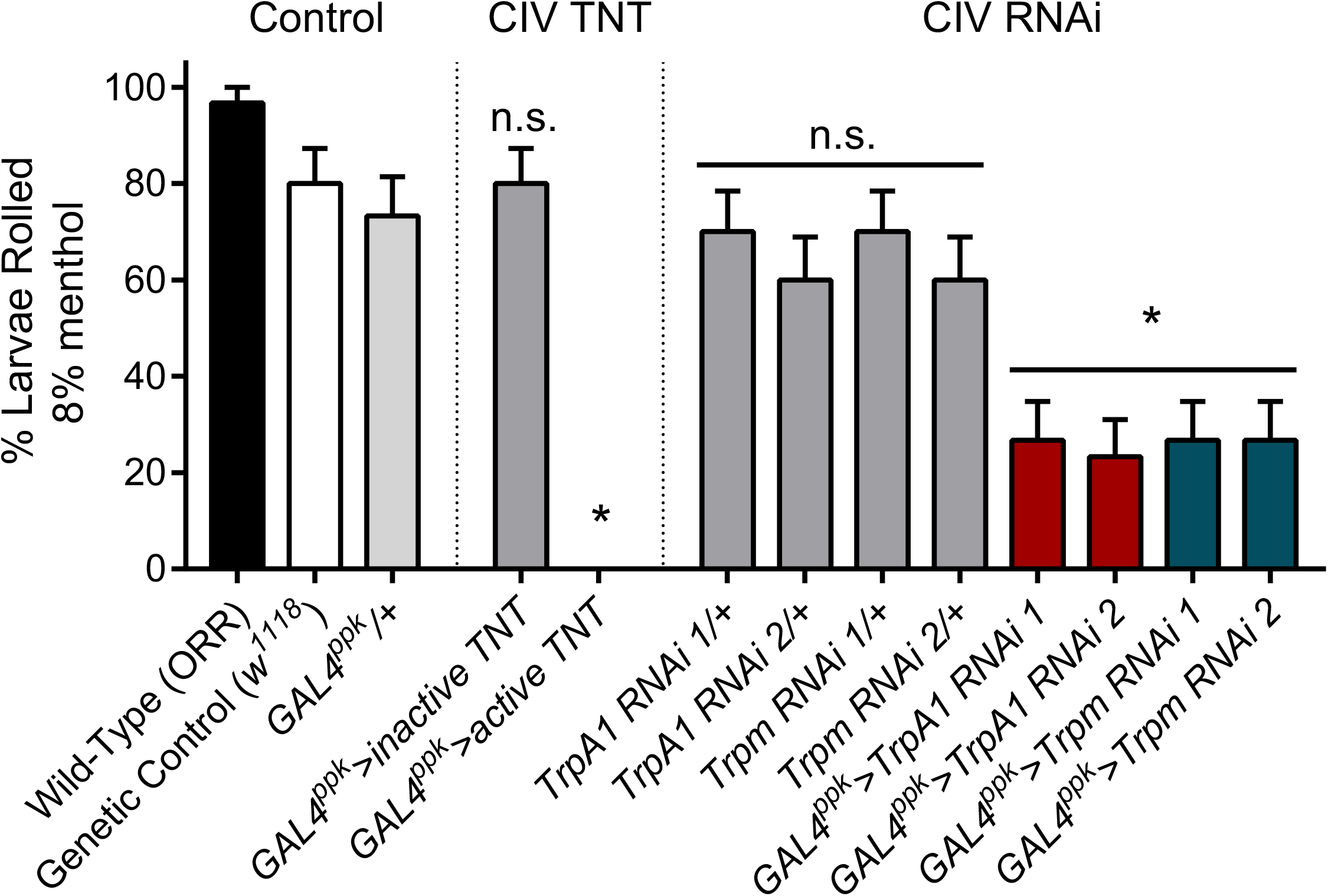
The TRP-dependent menthol response operates via CIV neurons. Silencing neural transmission in CIV neurons with active tetanus toxin (TNT) completely ablated the menthol response, while CIV-driven expression of inactive TNT did not significantly affect the rolling response. CIV-specific, RNAi-mediated knockdown using two independent transgenes for *TrpA1* and *Trpm* resulted in fewer animals rolling in response to menthol, as compared to control (*TrpA1* RNAi 1, p=0.0032; *TrpA1* RNAi 2, p=0.0018; *Trpm* RNAi 1, p=0.0032; and *Trpm* RNAi 2, p=0.0032). Proportions represented as % ± SEP. n=30 for each condition.

In light of TRP mutant and CIV TNT phenotypes, we hypothesized that menthol-evoked behaviour may be dependent on TrpA1 and/or Trpm function in CIV nociceptors. We previously demonstrated that *TrpA1* and *Trpm* transcripts are detectibly expressed in CIV nociceptors ([80], GEO: GSE46154). Consistent with these data, CIV-targeted, RNAi-mediated knockdown of *Trpm* or *TrpA1* strongly inhibited menthol-evoked rolling (**Figure 4**), supporting a functional role for these TRP channels in CIV-mediated menthol sensing.

### Residues critical to menthol sensing were likely present in ancestral bilaterian channels

A number of specific amino acid residues have been associated with menthol sensitivity in mammalian TRPM [12, 14, 16, 19, 20] and TRPA [31] channels. Phylogenetic and sequence analyses were performed in order to assess how well conserved these residues are across taxa, and to infer their evolutionary history. The amino acid sequences for 69 TRPM channels and 56 TRPA channels (**Table S1**) were used to generate phylogenetic trees by both Bayesian and maximum likelihood approaches. Trees were constructed using previously and newly characterized TRP sequences across a variety of metazoan species, with a chaonoflagellate (*Monosiga brevicollis*) outgroup (**Figure 5A**). Both methods produced trees with largely consistent topologies (**Figures S1-S4**).

**Figure 5.**
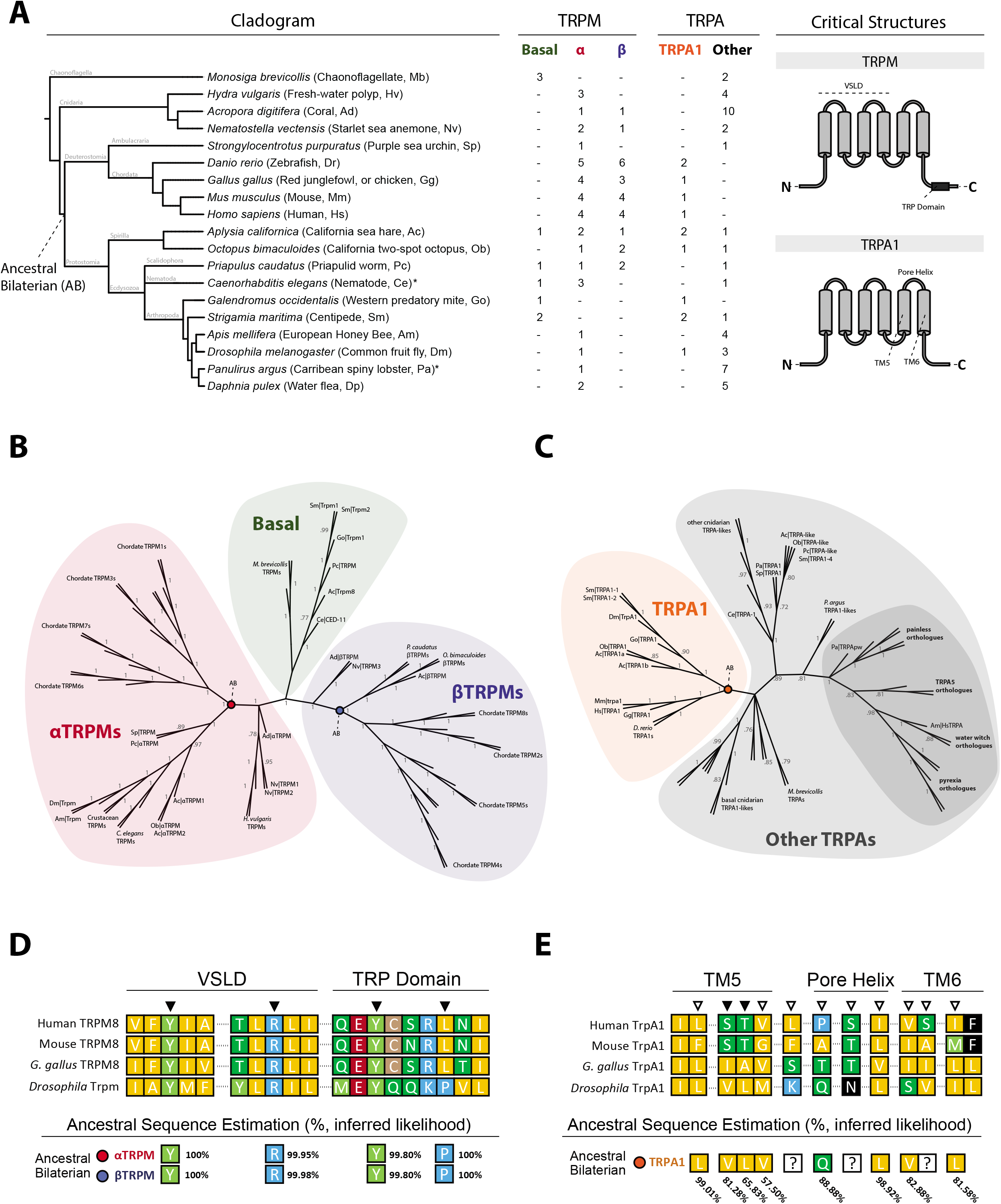
The evolutionary history of TRPM and TRPA channels, and of residues critical to menthol sensing. **(A)** (left) Cladogram of species used in analysis. Asterisk (*) indicates that one or more previously known sequences were discarded due to extreme divergence, as described in the Materials & Methods and Supplementary Materials. All sequence accession numbers are available in the Supplementary Materials. (right) Domains previously identified as critical to menthol sensing. **(B)** Bayesian consensus tree for TRPM channels. Posterior probabilities are indicated for each internal branch. Branches with a posterior probability <0.70 were collapsed. For aesthetic purposes, branch lengths were ignored when generating this figure. Trees with branch lengths to scale are available in the Supplementary Materials. Green, basal clade; red, αTRPM clade; blue, βTRPM clade. **(C)** Bayesian consensus tree for TRPA channels, as in B. Orange, TRPA1 clade; grey, other TRPAs; dark grey, clade previously described as “basal.” **(D)** Alignment for select TRPM sequences and ancestral sequence estimations. Black arrows indicate previously identified critical residue site. % values represent probability of amino acid at the indicated site, at the colour-coded node. **(E)** Alignment for select TRPA sequences and ancestral sequence estimations, as in D. White arrows indicate residues associated with species-specific menthol responses, in mammals. Question mark (?) indicates that no single amino acid was predicted with >50% inferred likelihood.

It has been previously noted that several species express TRPM-like channels which cluster independent of, or differ greatly from, other chordate and/or insect TRPMs [28, 52, 81]. In accordance with this, the consensus TRPM phylogeny shows a group of protostome TRPMs located basally, near chaonoflagellate TRPMs (**Figure 5B, green**). Previously published phylogenies have shown most arthropod TRPMs to be most closely related to chordate TRPMs 1, 3, 6, and 7, yet this relationship has not yet been formally discussed [25, 28]. These trees demonstrate that non-basal TRPM channels group into two monophyletic clades (designated αTRPM and βTRPM), which collectively constitute all other analyzed TRPM channels (**Figure 5B, red and blue**). Each clade contains both protostome and deuterostome TRPMs and is rooted in cnidarian TRPMs, indicating that these two clades may have existed prior to the protostome-deuterostome split. The topology of these trees is consistent with a hypothesis that at least three distinct TRPM channels (designated basal TRPM, αTRPM, and βTRPM) predate the last common bilaterian ancestor, Urbilateria. Further, this hypothesis is compatible with previously formulated hypotheses regarding the independent diversification of chordate TRPM channels [25, 28, 32].

As both α- and βTRPMs have been implicated in menthol sensing, ancestral sequence predictions were generated for ancestral bilaterian α- and βTRPM. Four TRPM8 residues – Y745, R842, Y1005, and L1009 (**Figure 5D, black arrows**) – have been identified as critical to vertebrate menthol sensing [20]. Three of these residues, Y745, R842, and Y1005, are conserved in *Drosophila*, and ancestral sequence predictions suggest with high confidence that they are conserved from a common ancestral bilaterian sequence (**Figure 5D, bottom**). The only site which differs from chordate TRPM8 is a predicted ancestral proline in place of L1009; proline, however, is more common at this site across TRPM channels, including in *Drosophila*, and previous work has shown that L1009P substitution in mouse TRPM8 does not affect menthol sensitivity [12].

For TRPAs, both previously characterized clades – TRPA1 and the “basal” clade (**Figure 5C, orange and dark grey**) – formed as expected, and conformed to other published topologies [7, 27, 28, 32]. Although the term “basal” has been used to describe other TRPAs, Peng, Shi, and Kadowaki have suggested that the TRPA1 clade is the most ancestral [28]. This analysis does not clearly support nor undermine this hypothesis. What is clear, however, is that the TRPA1 clade predates the protostome-deuterostome split.

Previous work has shown that the TRPA1 pore region is important to mammalian menthol sensitivity. Serine and threonine residues in TM5 are thought to be most critical to menthol sensitivity (**Figure 5E, black arrows**), and a variety of others (**Figure 5E, white arrows**) are associated with species-specific differences in menthol sensitivity [31]. These regions are generally poorly conserved between mammals and other species (**Figure 5E, top**), including other chordates (e.g., *Gallus gallus*). In contrast to TRPM conservation, fewer critical TRPA1 residues appear to be consistently conserved from a common bilaterian ancestor, and there is substantially less certainty as to the identity of these ancestral sequences (**Figure 5E, bottom**).

## Discussion

### TRP-dependent transduction and menthol sensing in *Drosophila*

We have demonstrated that *Drosophila* larvae execute a dose-dependent, menthol-evoked, aversive rolling response consistent with the nocifensive behaviour displayed in response to noxious heat and mechanical insult. Cellularly, menthol exposure activates CIV nociceptors, and blocking synaptic transmission in these neurons inhibits menthol-evoked rolling. Molecularly, TrpA1 and Trpm are required for menthol-induced aversive behaviour, and genetically interact in this capacity.

Curiously, it has been previously reported that *Drosophila* TrpA1 does not directly gate in response to menthol [31]. The study in question, however, used lower concentrations of menthol applied to channels expressed *in vitro*. These sorts of discrepancies are not uncommon in studies of other TRPA1 modalities, and there are similar conflicting reports concerning TRPA1’s role in cold nociception: some groups report that TRPA1 acts as a direct cold sensor, others state it responds to an indirect, cold-associated factor (reactive oxygen species and/or intracellular Ca^2+^), while others claim it responds only to innocuous cooling, or does not respond to cold at all [47, 82–86]; some isoforms of TRPA1 have been shown to directly gate in response to noxious high temperatures, yet high temperature nociception can be rescued by heat-insensitive isoforms [37]; and there is still debate as to whether or not TRPA1 functions similarly *in vivo* and *in vitro* [46]. Further, TRPA1s are differentially activated and/or inhibited by menthol, across species, across concentrations [41, 47]. It is therefore unsurprising, if no less puzzling, that *Drosophila TrpA1* is required for menthol detection *in vivo*, yet the channel does not gate in response to relatively low concentrations of menthol *in vitro*. Instead, these and other findings suggest that we may not fully understand how TRPA1 functions, and that it may not be a simple sensor which directly responds to an incredibly wide variety of stimuli.

Moreover, although menthol and TRPA/TRPM interactions are most frequently studied, menthol is likely to be very promiscuous with respect to which channels it activates. Reports suggest that menthol activates, potentiates, or inhibits voltage-gated Na^+^ channels, L-type voltage-gated Ca^2+^ channels, the cystic fibrosis transmembrane conductance regulator, mouse TRPV3, TRPL, a wide variety of Ca^2+^-dependent channels, as well as GABA, serotonin, and acetylcholine receptors [7, 87–91]. Another notable player is the hymenopteran specific TRPA (HsTRPA), which is more closely related to painless than to TRPA1 (**Figure 5C, dark grey**). HsTRPA1 thermal sensitivity is depressed by application of menthol [7]. Given that *painless* mutants are hypersensitive to menthol (**Figure 2C**), the so-called basal TRPAs may be complex mediators of chemo-sensitivity, rather than simple sensors. Currently, the menthol-specific ligand-binding and gating mechanisms of most receptors are not as well understood as those of chordate TRPA1 and TRPM8. However, these mechanisms are no doubt important, perhaps to species-specific behavioral responses, or to menthol’s purported analgesic effect [2–4, 88]. Future studies will need to investigate not only how other channels function in menthol sensing, but how they might interact with channels with incompletely understood activation properties (*e.g*., TRPA1), as these interactions may explain differences in menthol-evoked activity *in vivo* and *in vitro*.

### TRP channels and the evolution of chemical nociception

Phylogenetic analyses have revealed that 3 TRPM clades – designated basal, αTRPM, and βTRPM – and the TRPA1 clade, likely predate the protostome-deuterostome split. Further, many TRPM and TRPA1 residues critical to menthol sensing are conserved from ancestral bilaterian channels (although more variably so for TRPA1s). That residues critical to menthol sensing may predate the protostome-deuterostome split influences how we may understand the evolution of menthol production and avoidance. There is some evidence that menthol may be lethal to several insect species [10, 11], including *D. melanogaster* [9]; as such, the ability to sense and respond to menthol may be adaptive. Given that there is considerable latency to roll in response to menthol (**Figure 2D**), it is unclear if the aversive rolling behaviour is protective. Menthol sensing may be more important for adult females, which rather carefully select egg-laying sites and show aversion toward mentholated food [5]. Yet residues critical to menthol sensitivity substantially predate the emergence of Lamiaceae, the family of angiosperms that most notably produce menthol [92, 93]. This appears consistent with early plants evolving menthol (or more broadly, terpene) production in order to repel animals, rather than early animal TRPs evolving menthol sensitivity in order to avoid certain plants. It is possible that animal TRP channels maintained menthol sensitivity since well before terrestrial animals encountered menthol producers, much like how mammalian TRP channels are sensitive to icilin, which is not known to be naturally occurring [1, 16, 84, 94]. Pinpointing the exact origins of these abilities requires additional studies, in particular studies involving more basal (*e.g*. Xenacoelomorpha) and non-bilaterian species. Yet, it is conceivable that the functional capacity of these channels evolved in, or prior to, a common bilaterian ancestor, which inhabited a very different environment than extant terrestrial animals [95].

The structure of the urbilaterian nervous system is still under considerable debate [96–100]. However, given that TRP channels function in similar ways, in similar subsets of neurons, across taxa, it seems increasingly likely that ancestral TRP channels were expressed in some form of urbilaterian neural tissues, and that these tissues responded to external cues. Still, the function of conserved, menthol-associated residues in ancestral TRPs remains mysterious. However, while menthol production appears to be restricted to plants, terpene production (and more broadly, volatile organic compound production) is far more widespread. For example, volatile organic producers include plants, insects, protists, bacteria, and fungi [101], which have been hypothesized to utilize volatile organic compounds in forms of microbial communication [102]. With this in mind, and concerning terpene sensing by early metazoan TRPs, one plausible hypothesis is that TRP-terpene sensitivity emerged as an early method of communication. Alternatively (or perhaps as sub-hypotheses), TRP-terpene sensing may have been employed as a mechanism by which to identify appropriate food sources, or to signal potential danger, thereby avoiding damage from predation or infection – perhaps an early form of early nociception.

## Conclusion

Herein, we have described cellular and molecular mechanisms by which *Drosophila melanogaster* responds to menthol. Phylogenetic analyses revealed that extant TRPM channels are descended from 3 ancestral clades (basal, αTRPM, and βTRPM), and that several residues critical to menthol sensing are conserved from common TRPM and TRPA1 ancestors. Collectively, these findings suggest that bilaterian menthol sensing has its origins in a common ancestor, and more broadly, contribute to a body of evidence suggesting that the mechanisms underlying TRP function and chemical nociception are ancient and highly conserved.

## Acknowledgements

We would like to thank the Bloomington *Drosophila* Stock Center, as well as Drs Michael J. Galko, W. Dan Tracey, and Kartik Venkatachalam for providing *Drosophila* strains. This work was supported by the National Institute of Neurological Disorders and Stroke (NINDS) (R01NS086082; DNC); the National Institute of General Medical Sciences (NIGMS) IMSD (R25GM109442-01A1; TRG and DNC); Georgia State University (GSU) Brains & Behavior Seed Grant (DNC); GSU Brains & Behavior Fellowships (NJH and JML); and the Kenneth W. and Georganne F. Honeycutt Fellowship (NJH).

## Authors’ Contributions

Conceptualization and experimental design, NJH, JML, AS, and DNC; genetics, NJH and JML; behaviour assays, NJH, JML, MNB; CaMPARI microscopy, NJH and TRG; electrophysiology, AS; phylogenetic and sequence analyses, NJH; statistical analyses, NJH; wrote the original manuscript, NJH; critically analysed and edited the manuscript, NJH, JML, AS, TRG, MNB and DNC; approved the final draft, NJH, JML, AS, TRG, MNB, and DNC; funding acquisition, NJH and DNC.

