## Supplementary material for "*Drosophila* menthol sensitivity and the Precambrian origins of TRP-dependent chemosensation": Table S1 & Figures S1-S4

**Table of Contents** *Page*


| | |
|---|---|
| Table S1: Protein sequence information | 2 |
| Figure S1: TRPM maximum likelihood tree | 3 |
| Figure S2: TRPM Bayesian tree | 4 |
| Figure S3: TRPA maximum likelihood tree | 5 |
| Figure S4: TRPA Bayesian tree | 6 |
| Supplemental Materials References | 7 |
| Movie S1: Menthol evokes rolling behaviour | external |
| Movie S2: Vehicle (DMSO) does not evoke rolling | external |
| Movie S3: Icilin does not evoke rolling | external |

### Table S1

| Transient Receptor Potential, subfamily M | | | |
|---|---|---|---|
| Species | Protein | Source/Database | Accession/ID |
| **Choanoflagellate** | | | |
| *Monosiga brevicollis* | Mb\|TRPM1 | JGI Genome | Monbr1\|27446 |
| | Mb\|TRPM2 | JGI Genome | Monbr1\|29245 |
| | Mb\|TRPM3 | JGI Genome | Monbr1\|34057 |
| **Cnidaria** | | | |
| *Acropora digitifera* | Ad\|αTRPM | OIST Marine Genomics Unit | v2a.06725.t1 |
| | Ad\|βTRPM | OIST Marine Genomics Unit | v2a.10270.t1 |
| *Hydra vulgaris* | Hv\|TRPM1 | NCBI | XP_002168052.2 |
| | Hv\|TRPM2 | NCBI | XP_002167966.2 |
| | Hv\|TRPM3 | NCBI | XP_002163434.2 |
| *Nematostella vectensis* | Nv\|TRPM1 | JGI Genome | Nemve1\|102167 |
| | Nv\|TRPM2 | JGI Genome | Nemve1\|85952 |
| | NV\|TRPM3 | JGI Genome | Nemve1\|248535 |
| **Deuterostomia** | | | |
| *Danio rerio* | Dr\|TRPM1a | UniProt | AGS55979.1 |
| | Dr\|TRPM1b | UniProt | AGS55980.1 |
| | Dr\|TRPM2 | UniProt | AGS55981.1 |
| | Dr\|TRPM3 | UniProt | AGS55982.1 |
| | Dr\|TRPM4a | UniProt | AGS55983.1 |
| | Dr\|TRPM4b1 | UniProt | AGS55984.1 |
| | Dr\|TRPM4b2 | UniProt | AGS55985.1 |
| | Dr\|TRPM4b3 | UniProt | AGS55986.1 |
| | Dr\|TRPM5 | UniProt | AGS55987.1 |
| | Dr\|TRPM6 | UniProt | AGS55988.1 |
| | Dr\|TRPM7 | UniProt | AGS55989.1 |
| *Gallus gallus* | Gg\|TRPM1 | UniProt | F1NCK1 |
| | Gg\|TRPM2 | UniProt | F1NGK6 |
| | Gg\|TRPM3 | UniProt | F1NJZ7 |
| | Gg\|TRPM5 | UniProt | E1BXA4 |
| | Gg\|TRPM6 | UniProt | E1C5L2 |
| | Gg\|TRPM7 | UniProt | E1BYV3 |
| | Gg\|TRPM8 | UniProt | E1BWL7 |
| *Homo sapiens* | Hs\|TRPM1 | NCBI CCDS | 10024.2 |
| | Hs\|TRPM2 | NCBI CCDS | 13710.1 |
| | Hs\|TRPM3 | NCBI CCDS | 6634.1 |
| | Hs\|TRPM4 | NCBI CCDS | 33073.1 |
| | Hs\|TRPM5 | NCBI CCDS | 31340.1 |
| | Hs\|TRPM6 | NCBI CCDS | 6647.1 |
| | Hs\|TRPM7 | NCBI CCDS | 42035.1 |
| | Hs\|TRPM8 | NCBI CCDS | 33407.1 |
| *Mus musculus* | Mm\|Trpm1 | NCBI CCDS | 21332.2 |
| | Mm\|Trpm2 | NCBI CCDS | 48611.1 |
| | Mm\|Trpm3 | NCBI CCDS | 29702.1 |
| | Mm\|Trpm4 | NCBI CCDS | 52245.1 |
| | Mm\|Trpm5 | NCBI CCDS | 52462.1 |
| | Mm\|Trpm6 | NCBI CCDS | 29691.1 |
| | Mm\|Trpm7 | NCBI CCDS | 16689.1 |
| | Mm\|Trpm8 | NCBI CCDS | 48316.1 |
| *Strongylocentrotus purpuratus* | Sp\|TRPM | EchinoBase | SPU_014937-tr |
| **Protostomia** | | | |
| *Apis mellifera* | Am\|Trpm | NCBI | XP_395849.4 |
| *Aplysia californica* | Ac\|TRPM-like | NCBI | XP_012937083.1 |
| | Ac\|αTRPM1 | NCBI | XP_012936450.1 |
| | Ac\|αTRPM2 | NCBI | XP_012936451.1 |
| | Ac\|βTRPM | NCBI | XP_012940727.1 |
| *Caenorhabditis elegans* | Ce\|gon-2 | WormBase | CE30390 |
| | Ce\|gtl-1 | WormBase | CE33754 |
| | Ce\|gtl-2 | WormBase | CE40563 |
| | Ce\|ced-11 | WormBase | CE00409 |
| *Daphnia pulex* | Dp\|TRPM1 | JGI Genome | Dappu\|51380 |
| | Dp\|TRPM2 | JGI Genome | Dappu\|51222 |
| *Drosophila melanogaster* | Dm\|Trpm | FlyBase | FBgn0265194 |
| *Galendromus occidentalis* | Go\|TRPM1 | NCBI | XP_003744025.1 |
| *Octopus bimaculoides* | Ob\|αTRPM | NCBI | XP_014787598.1 |
| | Ob\|βTRPM1 | NCBI | XP_014775201.1 |
| | Ob\|βTRPM2 | NCBI | XP_014774531.1 |
| *Panulirus argus* | Pa\|TRPM | (Kozma et al., 2018) | - |
| | Pa\|TRPMm* | (Kozma et al., 2018) | - |
| *Priapulus caudatus* | Pc\|TRPM-like | NCBI | XP_014670278.1 |
| | Pc\|αTRPM | NCBI | XP_014671784.1 |
| | Pc\|βTRPM1 | NCBI | XP_014666350.1 |
| | Pc\|βTRPM2 | NCBI | XP_014669422.1 |
| *Strigamia maritima* | Sm\|Trpm1 | EnsemblMetazoa | SMAR010526-RA |
| | Sm\|Trpm2 | EnsemblMetazoa | SMAR006477-RA |

| Transient Receptor Potential, subfamily A | | | |
|---|---|---|---|
| Species | Protein | Source/Database | Accession/ID |
| **Choanoflagellate** | | | |
| *Monosiga brevicollis* | Mb\|TRPA1-1 | JGI Genome | Monbr1\|22692 |
| | Mb\|TRPA1-2 | JGI Genome | Monbr1\|28830 |
| **Cnidaria** | | | |
| *Acropora digitifera* | Ad\|TRPA1-like1a | OIST Marine Genomics Unit | v2a.19411.t1 |
| | Ad\|TRPA1-like1b | OIST Marine Genomics Unit | v2a.12507.t1 |
| | Ad\|TRPA1-like2a | OIST Marine Genomics Unit | v2a.03973.t1 |
| | Ad\|TRPA1-like2b | OIST Marine Genomics Unit | v2a.12510.t1 |
| | Ad\|TRPA1-like2c | OIST Marine Genomics Unit | v2a.12511.t1 |
| | Ad\|TRPA1-like2d | OIST Marine Genomics Unit | v2a.12512.t1 |
| | Ad\|TRPA1-like2e | OIST Marine Genomics Unit | v2a.12513.ta |
| | Ad\|TRPA1-like3 | OIST Marine Genomics Unit | v2a.17511.t1 |
| | Ad\|TRPA-like1a | OIST Marine Genomics Unit | v2a.00690.t1 |
| | Ad\|TRPA-like1b | OIST Marine Genomics Unit | v2a.00691.t1 |
| *Hydra vulgaris* | Hv\|TRPA1-1 | NCBI | XP_002167191.2 |
| | Hv\|TRPA1-2 | NCBI | XP_002169693.2 |
| | Hv\|TRPA1-3 | NCBI | XP_002164531.2 |
| | Hv\|TRPA1-4 | NCBI | XP_002159822.2 |
| *Nematostella vectensis* | Nv\|TRPA1-1 | JGI Genome | Nemve1\|224166 |
| | Nv\|TRPA1-2 | JGI Genome | Nemve1\|217023 |
| **Deuterostomia** | | | |
| *Danio rerio* | Dr\|TRPA1a | UniProt | Q5UM15 |
| | Dr\|TRPA1b | UniProt | Q5UM14 |
| *Gallus gallus* | Gg\|TRPA1 | UniProt | W8VTH6 |
| *Homo sapiens* | Hs\|TRPA1 | NCBI CCDS | 34908.1 |
| *Mus musculus* | Mm\|TRPA1 | NCBI CCDS | 35516.1 |
| *Strongylocentrotus purpuratus* | Sp\|TRPA1 | EchinoBase | SPU_010335 |
| **Protostomia** | | | |
| *Apis mellifera* | Am\|Pain | NCBI | XP_001122160.2 |
| | Am\|TRPA5 | NCBI | XP_001122445.1 |
| | Am\|pyx | NCBI | XP_006570269.1 |
| | Am\|wtrw | NCBI | XP_395234.3 |
| | Am\|HsTRPA | NCBI | XP_395235.2 |
| *Aplysia californica* | Ac\|TRPA1-like | NCBI | XP_012939173.1 |
| | Ac\|TRPA1a | NCBI | XP_012939582.1 |
| | Ac\|TRPA1b | NCBI | XP_012937564.1 |
| *Caenorhabditis elegans* | Ce\|trpa-1 | WormBase | C29E6.2a.1 |
| | Ce\|trpa-2* | Wormbase | CE18081 |
| *Daphnia pulex* | Dp\|pyx1 | NCBI | EFX90080.1 |
| | Dp\|pyx2 | JGI Genome | Dappu1\|327376\| |
| | Dp\|pyx3 | JGI Genome | Dappu1\|329422 |
| | Dp\|pain | JGI Genome | Dappu1\|315097 |
| | Dp\|TRPA5 | JGI Genome | Dappu1\|318074 |
| *Drosophila melanogaster* | Dm\|TrpA1 | FlyBase | FBpp0304207 |
| | Dm\|pain | FlyBase | FBpp0072323 |
| | Dm\|pyx | FlyBase | FBpp0072423 |
| | Dm\|wtrw | FlyBase | FBpp0081250 |
| *Galendromus occidentalis* | Go\|TrpA1 | NCBI | XP_003748128.1 |
| *Octopus bimaculoides* | Ob\|TRPA-like | NCBI | XP_014772761.1 |
| | Ob\|TRPA1 | NCBI | XP_014785409.1 |
| *Panulirus argus* | Pa\|TRPApw | (Kozma et al., 2018) | - |
| | Pa\|pain1 | (Kozma et al., 2018) | - |
| | Pa\|pain2 | (Kozma et al., 2018) | - |
| | Pa\|TRPA1-like1 | (Kozma et al., 2018) | - |
| | Pa\|TRPA1-like2 | (Kozma et al., 2018) | - |
| | Pa\|TRPA5-1* | (Kozma et al., 2018) | - |
| | Pa\|TRPA5-2 | (Kozma et al., 2018) | - |
| | Pa\|TRPA-like1* | (Kozma et al., 2018) | - |
| | Pa\|TRPA1 | (Kozma et al., 2018) | - |
| *Priapulus caudatus* | Pc\|TRPA-like | NCBI | XP_014673121.1 |
| *Strigamia maritima* | Sm\|TRPA1-1 | EnsemblMetazoa | SMAR011250-RA |
| | Sm\|TRPA1-2 | EnsemblMetazoa | SMAR011251-RA |
| | Sm\|TRPA1-4 | EnsemblMetazoa | SMAR001905-RA |

**Table S1.** Name, source, and accession numbers for amino acid sequences used in phylogenetic analyses. Asterisk (*) indicates sequence was excluded due to extreme divergence, as described in Materials & Methods.

### Figure S1

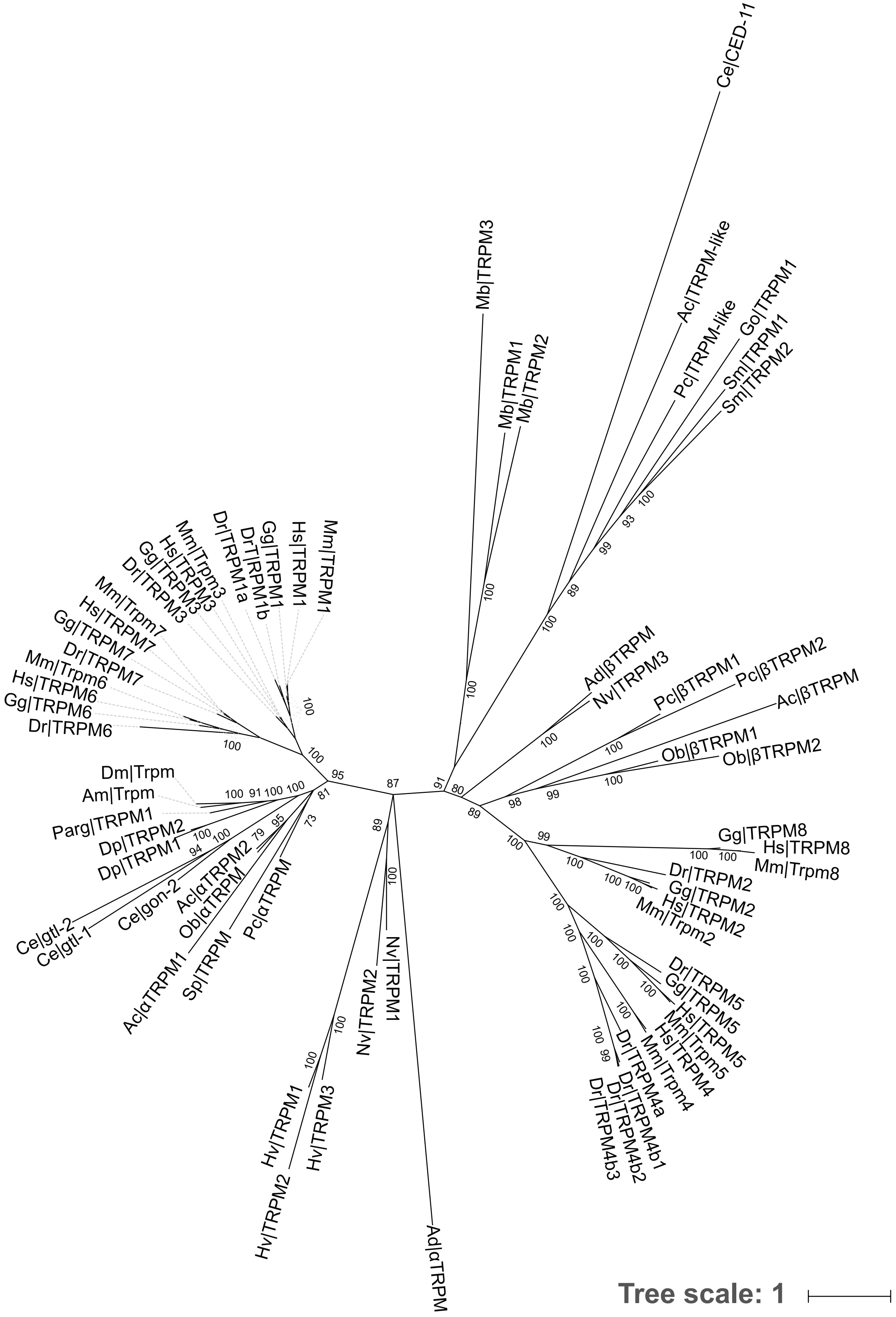


**Figure S1.** TRPM maximum likelihood tree. Node values indicate bootstrap confidence. Bootstraps <70% have been collapsed. Rooted in *M. brevicollis* TRPM equences.

### Figure S2

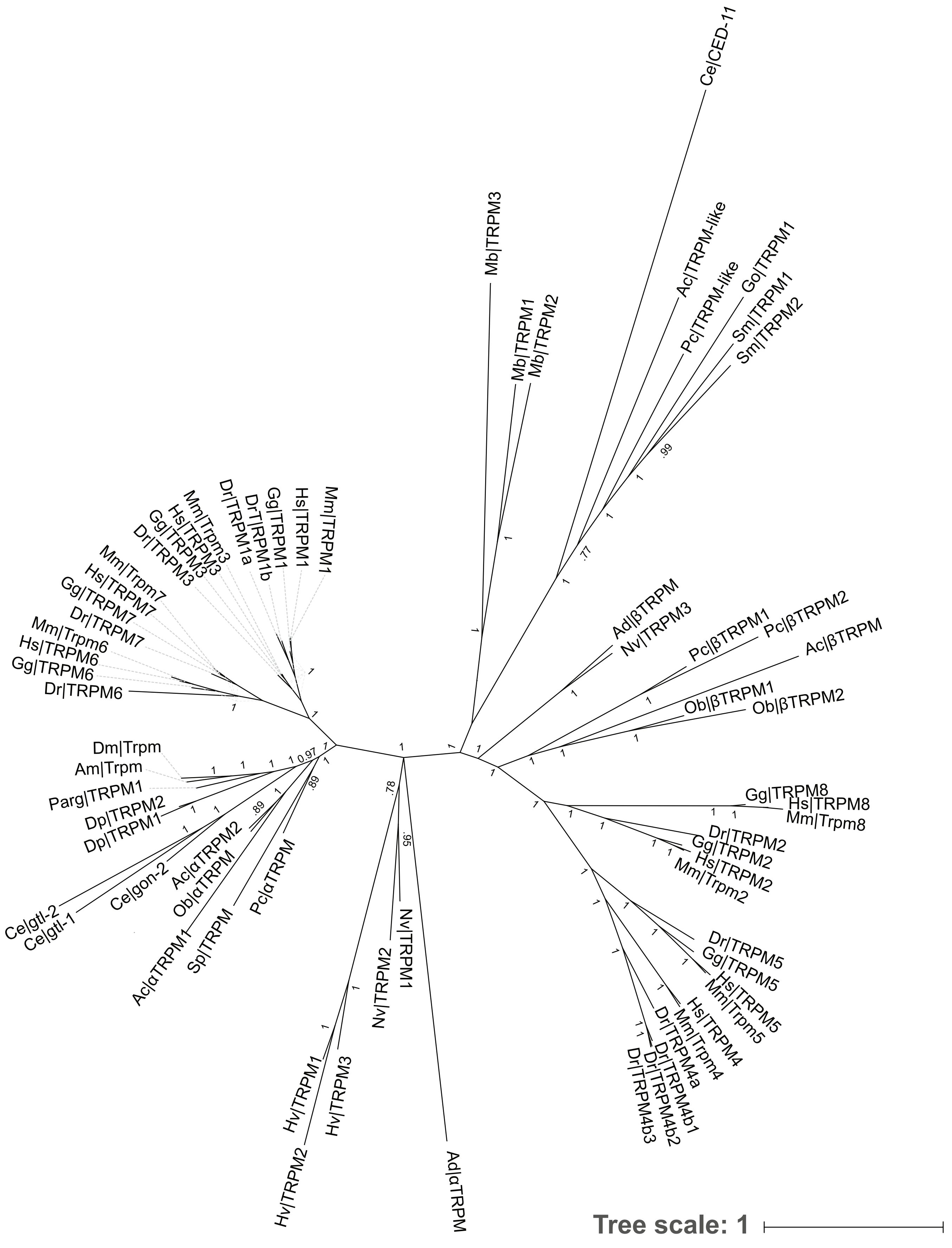


**Figure S2.** TRPM Bayesian tree. Node values are posterior probabilities for that branch. Posterior probabilities <0.70 have been collapsed. Rooted in *M. brevicollis* TRPM sequences.

### Figure S3

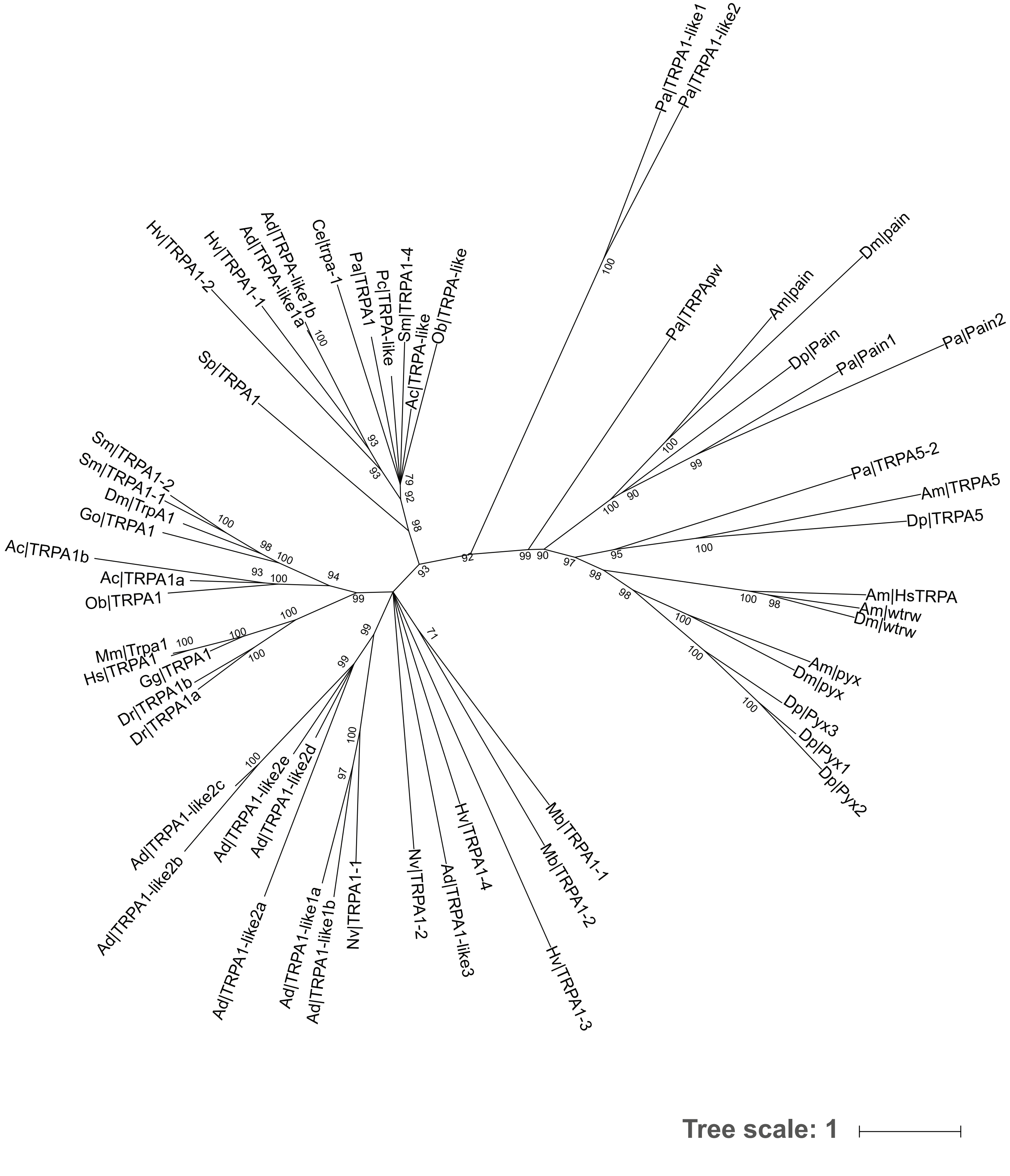


**Figure S3.** TRPA maximum likelihood tree. Node values indicate bootstrap confidence. Bootstraps <70% have been collapsed. Rooted in *M. brevicollis* TRPA sequences.

### Figure S4

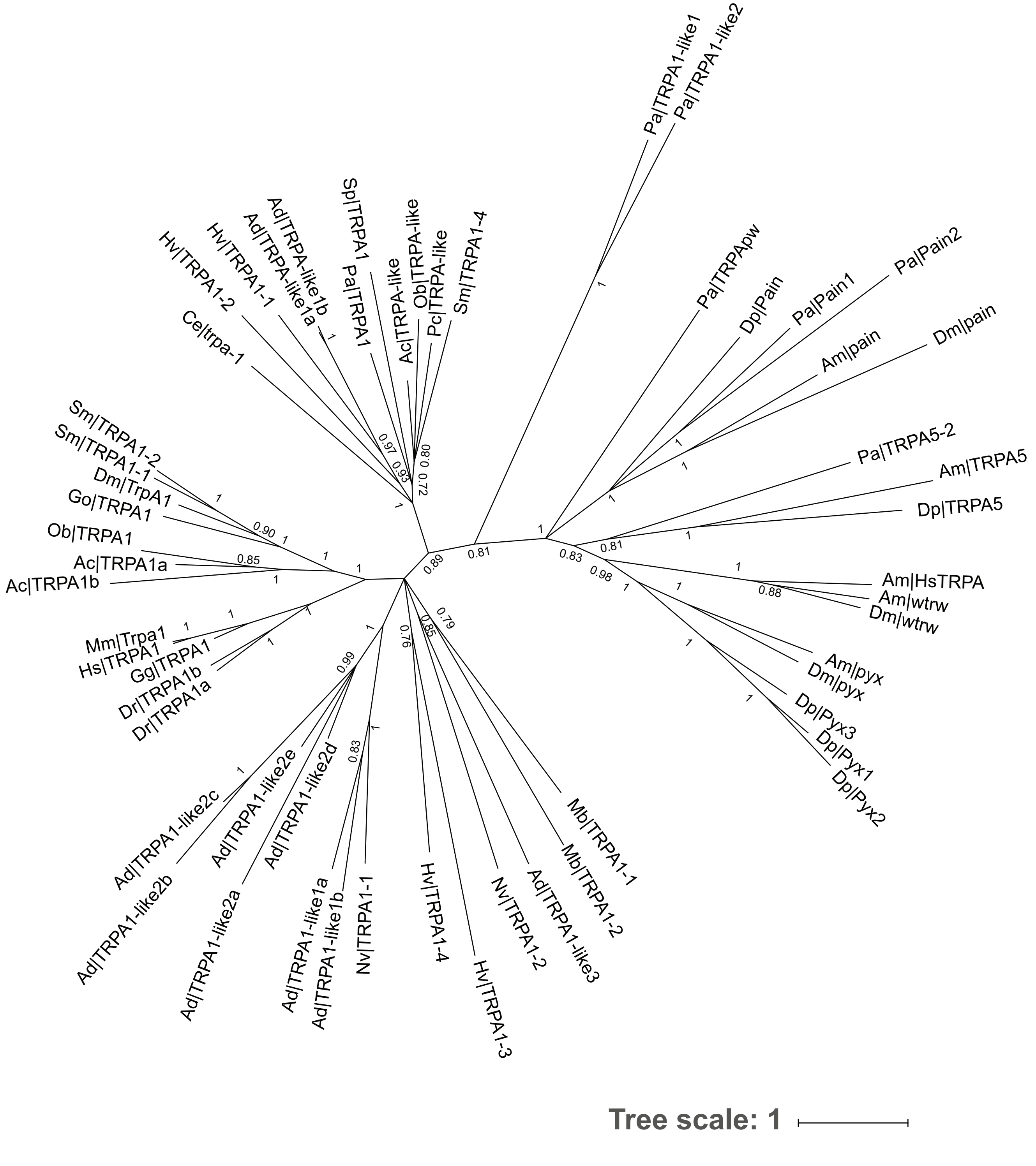


**Figure S4.** TRPA Bayesian tree. Node values are posterior probabilities for that branch. Posterior probabilities <0.70 have been collapsed. Rooted in *M. brevicollis* TRPA sequences.
